# Bacteria evolve macroscopic multicellularity via the canalization of phenotypically plastic cell clustering

**DOI:** 10.1101/2022.09.20.508687

**Authors:** Yashraj Chavhan, Sutirth Dey, Peter A. Lind

## Abstract

The evolutionary transition from unicellular to multicellular life was a key innovation in the history of life. Given scarce fossil evidence, experimental evolution has been an important tool to study the likely first step of this transition, namely the formation of undifferentiated cellular clusters. Although multicellularity first evolved in bacteria, the extant experimental evolution literature on this subject has primarily used eukaryotes. Moreover, it focuses on mutationally driven (and not environmentally induced) phenotypes. Here we show that both Gram-negative and Gram-positive bacteria exhibit phenotypically plastic (*i.e.*, environmentally induced) cell clustering. Under high salinity, they grow as elongated ~ 2 cm long clusters (not as individual planktonic cells). However, under habitual salinity, the clusters disintegrate and grow planktonically. We used experimental evolution with *Escherichia coli* to show that such clustering can be canalized successfully: the evolved bacteria inherently grow as macroscopic multicellular clusters, even without environmental induction. Highly parallel mutations in genes linked to cell wall assembly formed the genomic basis of canalized multicellularity. While the wildtype also showed cell shape plasticity across high versus low salinity, it was either canalized or reversed after evolution. Interestingly, a single mutation could canalize multicellularity by modulating plasticity at multiple levels of organization. Taken together, we show that phenotypic plasticity can prime bacteria for evolving undifferentiated macroscopic multicellularity.

## Introduction

The evolutionary shift from unicellular organisms to multicellular ones represents an important gateway towards innovation in the history of life^1^. This shift has conventionally been categorized as a ‘major evolutionary transition’ because it created a new level of biological organization that natural selection could act on^2^, which likely facilitated an unprecedented increase in biological complexity^3,4^. Here we focus on the evolution of the capacity to form undifferentiated cellular clusters, which was likely the first key step towards the evolution of multicellularity and has evolved independently in at least 25 distinct lineages across the tree of life^1,3,5^. Discerning the nuances of this evolutionary transition is inherently difficult because it occurred in deep past > 2 billion years ago. Most transitional forms have likely undergone extinction, and the scarcity of fossil evidence severely limits what can be gleaned about this transition. In the face of severely limited fossil evidence, experimental evolution has proven to be a very powerful tool in this regard as it can combine empirical rigor with diverse experimental designs to directly observe the unfolding of this transition in action^6–11^.

Most studies on the experimental evolution of multicellularity in ancestrally unicellular organisms have dealt with eukaryotes^6,7,9,11–15^. Unicellular fungi^6,9,10^ and algae^11,12,14^ have proven to be particularly useful model organisms in this context. Moreover, experimental evolution approaches for studying multicellularity have been extended to a ‘non-model’ ichthyosporean relative of animals^8^. Furthermore, a recent experimental evolution study has even succeeded in demonstrating the evolution of macroscopic multicellularity in yeast^9^. Interestingly, multicellularity has independently evolved in prokaryotes at least three different times in the history of life^3,16^. However, there have been very few prokaryotic experimental evolutional studies demonstrating the *de novo* evolution of multicellularity, and these studies have been largely restricted to mat formation in *Pseudomonads* ^17^. In contrast, a rich body of work has investigated the nuances of the *already* evolved prokaryotic multicellularity^18–20^. Some particularly striking examples include the fruiting bodies of myxobacteria^21^, filamentous growth with cellular differentiation in cyanobacteria^22^, and the complex hyphal networks of streptomycetes^23^. Thus, the scarcity of prokaryotic experimental evolution studies represents a key gap in the current understanding of the evolution of multicellularity, which we aim to address here.

Another important aspect of most experimental evolution studies on multicellularity is their focus on mutationally derived (not environmentally induced) multicellular phenotypes^8,9,13–15^. These studies have revealed that the mutations required to form undifferentiated multicellular clusters are relatively easily accessible in diverse unicellular eukaryotic taxa (REFs). Moreover, a wide variety of environmental conditions can selectively enrich such *de novo* mutations (e.g., predation^7,11,12^, diffusible stressful agents^24^, improved extracellular metabolism^13^, etc. (reviewed in Ref. 5). Interestingly, novel phenotypes like multicellular clusters can also be expressed in the absence of mutations: phenotypic plasticity, which enables a given genotype to express different phenotypes in different environments^25–27^, is the basis of the facultative multicellular phenotypes exhibited by diverse taxa. For example, phytoplankton^28^, cyanobacteria^29^, and *Pseudomonads*^30,31^ can facultatively form multicellular clusters in response to predation. Moreover, a recent study has shown that changes in environmental salinity can induce multicellular clustering in marine cyanobacteria^32^. Such prevalence of facultative cell clustering across diverse unicellular taxa suggests that phenotypic plasticity may be an important force in the evolution of multicellularity. This is because plasticity can facilitate biological innovation by allowing genes to be ‘followers’ in the evolution of new phenotypes^33,34^. Specifically, selection can act on plastic phenotypes and enrich mutations which can canalize their expression and make them constitutively expressed, even in the absence of environmental induction^35–37^. Indeed, such canalization has been demonstrated in the unicellular alga *Chlamydomonas reinhardtii*^11^ (also see Ref. 7), which facultatively forms microscopic clusters (palmelloids) comprising ~ 140 cells in the presence of rotifer predators. Becks and colleagues showed that a sustained exposure to rotifer predators for 6 months led to constitutive palmelloid development in *C. reinhardtii*^11^. However, apart from this study, no other experiment has demonstrated that phenotypic plasticity can facilitate the evolution of multicellularity. Two specific questions remain unanswered in this context: (1) Can phenotypic plasticity facilitate the evolution of *macroscopic* multicellularity (comprising large clusters with > 10^4^ cells)? (2) Can it do so in bacteria? Our study addresses both these questions empirically.

Here we show that phenotypic plasticity can facilitate the evolution of macroscopic multicellularity in bacteria by bypassing and avoiding the wait for mutational emergence of undifferentiated cluster formation. We demonstrate that phenotypically plastic cell clustering in ancestral genotypes can be rapidly canalized to efficiently form multicellular clusters even in the absence of the environmental induction. We elucidate that phenotypically plastic clustering is also manifested at the level individual cell shapes. Finally, we show that mutations in a small number of genes linked to the cell wall can canalize the ancestral phenotypic plasticity at multiple levels of organizations, ultimately leading to obligately multicellular bacterial life histories.

## Results

### Both Gram-negative and Gram-positive bacteria exhibit phenotypically plastic cell clustering

We observed that high salinity liquid environments can make both Gram-negative and Gram-positive bacteria grow primarily as elongated macroscopic clusters and not as turbid cultures of individual planktonic cells (Fig. 1). Specifically, we grew independent clonal cultures of *Escherichia coli* (Gram-negative) and *Staphylococcus aureus* (Gram-positive) in two distinct environments (Luria Bertani broth containing either 0.5% or 6% NaCl (w/vol)) in unshaken tubes at 37°C (see *Methods*). Henceforth, we refer to these two environments as “habitual salinity” and “high salinity”, respectively. These two bacterial species have putatively diverged from their common ancestor >3000 million years ago (see *Methods*). As expected, under habitual salinity, both *E. coli* and *S. aureus* showed planktonic turbid growth without any observable clustering (Fig. 1; Supplementary movies 1 and 2). In contrast, under high salinity, both *E. coli* and *S. aureus* grew predominantly as elongated clusters and not as planktonic cultures (Fig. 1; Supplementary movies 3 and 4). In both species, the clusters comprised > 10^5^ viable colony forming units (CFUs) and reached 2-3 cm in length when cultured in tubes containing 5 ml nutrient medium. Such clustering was phenotypically plastic (environmentally induced): when transferred to a habitual salinity environment, the clusters disintegrated into individual cells that grew planktonically (Fig. S1). High-resolution time-lapse videos of macroscopic cluster formation revealed that in static high salinity environments, both *E. coli* and *S. aureus* showed a combination of clonal and aggregative modes of multicellular growth (Supplementary movies 2 and 4). Put differently, the multicellular growth under high salinity was a consequence of bacterial cells staying together after division (clonal expansion) *and* previously unattached cells (or cellular clusters) adhering to each other (aggregative growth).

**Figure 1.**
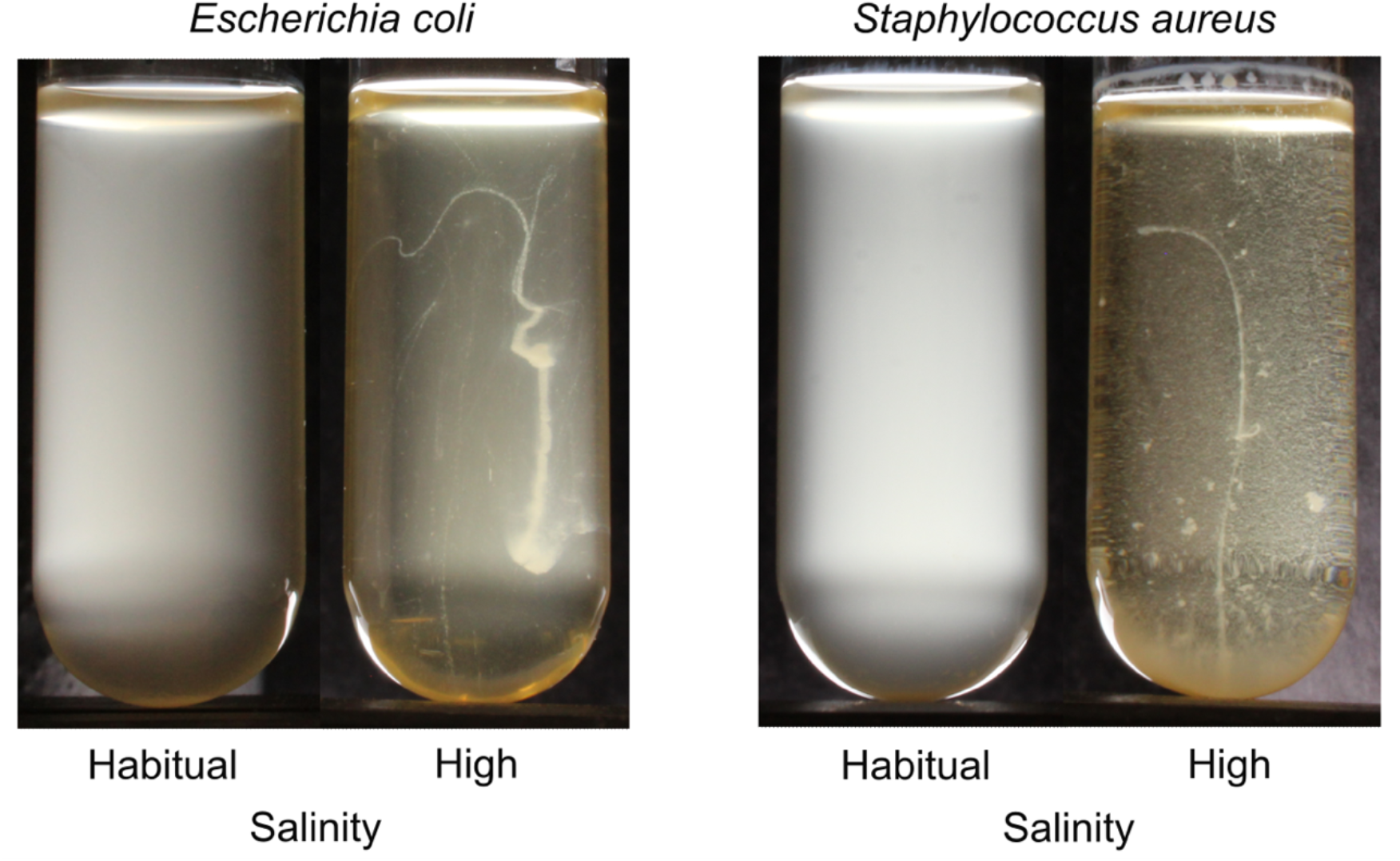
Both *E. coli* (Gram negative) and *S. aureus* (Gram positive) show the capacity to form phenotypically plastic elongated macroscopic cell clusters.

We further established that both *Citrobacter freundii* and *Pseudomonas aeruginosa* also exhibit such environmentally induced cell clustering suggesting that it is widespread in bacteria (Fig. S2). However, the formation of elongated clusters is not a physically inevitable outcome of bacterial growth under high salinity: the Gram-negative bacterium *Serratia marcescens* did not exhibit such phenotypic plasticity and grew as a turbid planktonic culture under both habitual and high salinity (Fig. S2). This led us to investigate if the phenotypically plastic bacterial clustering was itself an evolvable biological phenomenon.

Since the emergence of undifferentiated clusters is expected to be the first key step towards the evolution of multicellularity^1,3,5^, we hypothesized that phenotypically plastic cell clustering could facilitate the evolution of undifferentiated multicellularity in bacteria. Focusing on *Escherichia coli*, we set out to determine if the clustering induced by high salinity can be canalized into an obligately multicellular bacterial life history, even in the absence of environmental induction.

### Experimental evolution of simple multicellularity via the canalization of phenotypically plastic cell clustering

We established that *E. coli* could form phenotypically plastic macroscopic clusters not only in resting tubes but also in well-mixed environments where the culture tubes were shaken at ~180 rpm (Fig. S3). We hypothesized that resting and shaken high salinity environments should offer different selection pressures during evolution experiments: In resting cultures, oxygen supply depletes steeply from the air-liquid interface to tube’s floor. Hence, selection for increased clustering in resting cultures is likely to enrich mutants that cluster preferentially at the air-liquid interface^38^. In contrast, such oxygen availability gradients are much weaker in shaken tubes, where selection for greater clustering may not enrich interface inhabiting mutants. Building on these ecological contrasts, we used a single *E. coli* MG1655 colony to propagate two distinct experimental evolution lines (S (for Shaken) and R (for Resting)) to select for increased cell clustering in environments with progressively reducing salinity (Fig. 2a; see *Methods*). Propagating five replicate populations per line, we started the evolution experiment with media containing 6% NaCl (w/vol) and progressively reduced the salt concentration over 50 days (see *Methods*). Our selection protocol was designed to weed out planktonic bacteria growing outside clusters (Fig. 2a; see *Methods*). Unlike most other evolution experiments, here the phenotype of interest (macroscopic cluster formation) was already exhibited by the ancestor at the outset (induced by high salinity). We hypothesized that selection for clustering under progressively reduced salinity should enrich mutations that can make the clustering relatively less dependent on environmental induction. This expectation mirrors the “genes as followers” view of phenotypic evolution^34^. At the end of the evolution experiment, we tested if clones from the evolved populations were able to make macroscopic clusters in static habitual salinity environments (see *Methods*).

**Figure 2.**
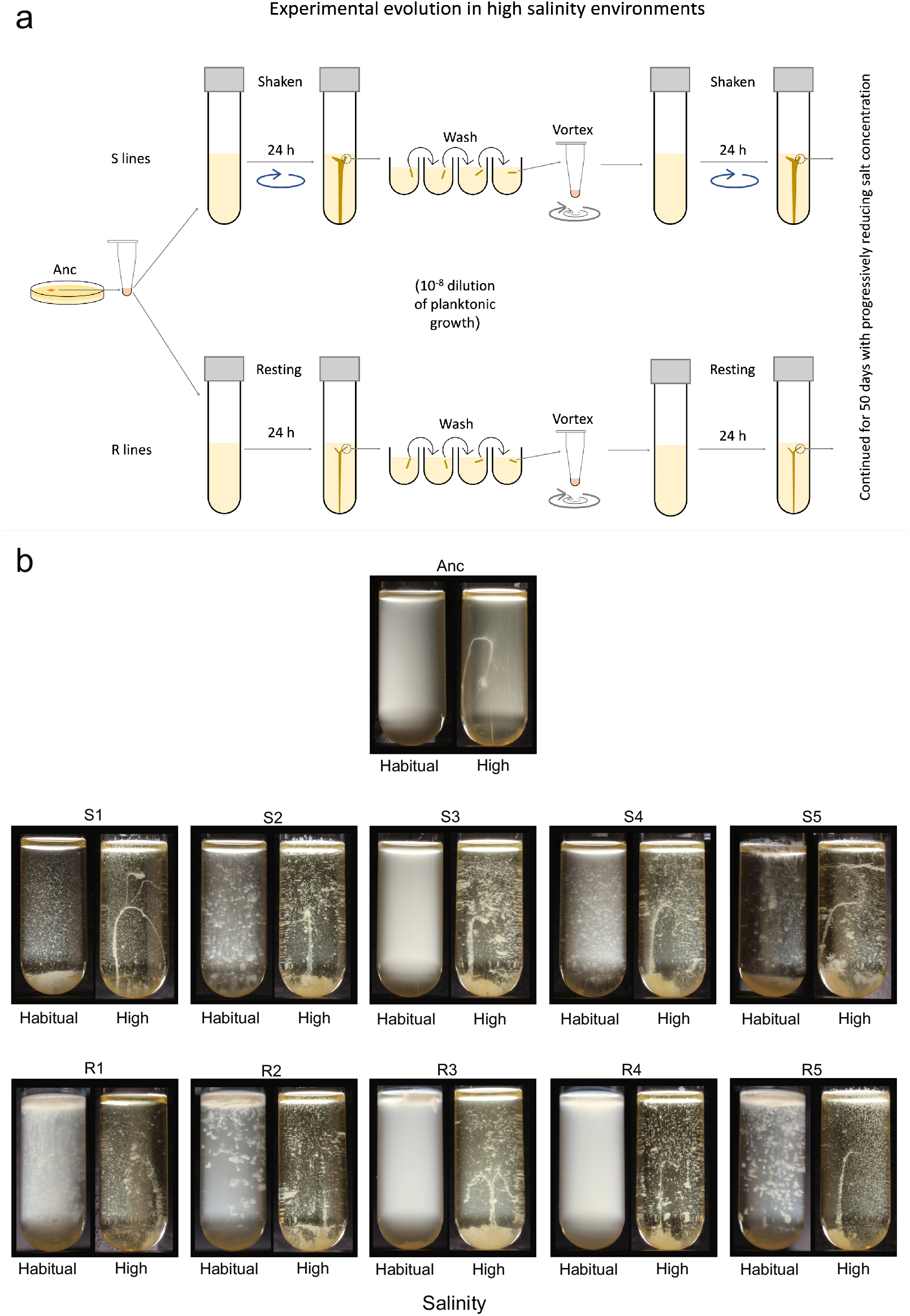
Experimental evolution of macroscopic multicellularity. (a) A schematic of our experimental evolution workflow. (b) Clonal phenotypes at the end of experimental evolution after growth under static conditions. Also see Supplementary movies 1 & 3 (for Anc), 5 & 7 (for the S clones), and 6 & 8 (for the R clones). In R1-R5, the habitual salinity tubes were externally perturbed at the end of the growth cycle to disrupt mats formed at the air-liquid interface and show cell clustering (see Fig. S5 for the unperturbed tubes).

Our evolution experiment successfully canalized the ancestrally plastic phenotype in most of the evolved lines (Fig. 2b; Supplementary movies 5 and 6). Specifically, clones representing 4 out of five S lines (S1, S2, S4, and S5) and 4 out of five R lines (R1, R2, R3, and R5) grew as macroscopic clusters even in the absence of environmental induction (Fig. 2b; Supplementary movies 5 and 6). Moreover, all five replicates of both S and R retained their ancestral ability to form elongated clusters in high salinity environments (Fig. 2b; Supplementary movies 7 and 8). Furthermore, under high salinity, both S and R showed significantly greater CFUs within their clusters as compared to the ancestor, suggesting an increase in the carrying capacity during selection (single sample t-tests against the ancestor: *P* = 0.0181 (for S); *P* = 0.021 (for R)); Fig. S4). Interestingly, the macroscopic clusters formed under habitual salinity were not elongated: S1, S2, S4, and S5 made a large number of macroscopic clusters that sank upon rapidly growing in size (Supplementary movie 5). The habitual salinity environment offers a weaker buoyant force than the high salinity environment; this could explain why the macroscopic clusters formed by S1, S2, S4, and S5 under habitual salinity were not elongated like the clusters formed by these clones under high salinity. In contrast to the S clones, R1, R2, R3, and R5 each formed a single mat (~1 mm thick) at the air-liquid interface under habitual salinity (Fig. S5; Supplementary movie 6). Moreover, the R1, R2, and R5 mats remained intact throughout the growth phase (Fig. S5); these mats disintegrated and sank only upon external perturbation, as shown in Fig. 2b. Thus, selection for clustering without environmental induction in resting tubes indeed enriched mutants that preferentially grew at the air-liquid interface, as we had hypothesized initially.

Since S1, S2, S4, S5, R1, R2, R3, and R5 formed multicellular clusters even in the absence of environmental induction, we conclude that they successfully evolved the first step towards multicellularity which demands that cells *inherently* grow as clusters. Interestingly, our selection protocol made the bacterial clusters undergo an artificially imposed life cycle where a small piece of the cluster in question (which was disintegrated by vigorous vertexing and then transferred into fresh media) gave rise to a new (larger) cluster. This motivated us to test if the clusters also qualify biological units that could spontaneously complete a life cycle consisting at least one multicellular stage^39–41^. To this end, we cultured an S clone (S5) in an arena where the bacteria could access fresh nutrients without being artificially transferred using a pipette (see *Methods*). We found that the bacteria successfully completed a life cycle where the old clusters gave rise to new clusters after accessing fresh nutrient medium, in both habitual and high salinity environments (Supplementary movie 9).

Taken together, the canalization of ancestrally plastic cell clustering led to the evolution of undifferentiated multicellularity in our experiments, which enabled bacteria to grow inherently as multicellular units, even in the absence of environmental induction. Having investigated phenotypic plasticity and its canalization at the level of *collectives* of cells (clusters), we turned our attention to the effects of selection on phenotypes at the level of *individual* cells.

### Phenotypic plasticity and its evolution at the cellular level

We performed both brightfield and fluorescence microscopy on the ancestral and evolved clones to determine if and how macroscopic cluster formation corresponded to changes in the cell shape (see *Methods*). We found that *E. coli* shows stark phenotypic plasticity in cell shape between habitual and high salinity environments (Fig. 3). Specifically, whereas the ancestral genotype showed its characteristic rod shape under habitual salinity, its cells became spherical under high salinity (Fig. 3). Surprisingly, we found that all the evolved lines lost their spherical cell shape under high salinity and their cells became elongated (Fig. 3). We quantitatively analyzed these cellular morphological changes using two distinct metrics (see *Methods*).

**Figure 3.**
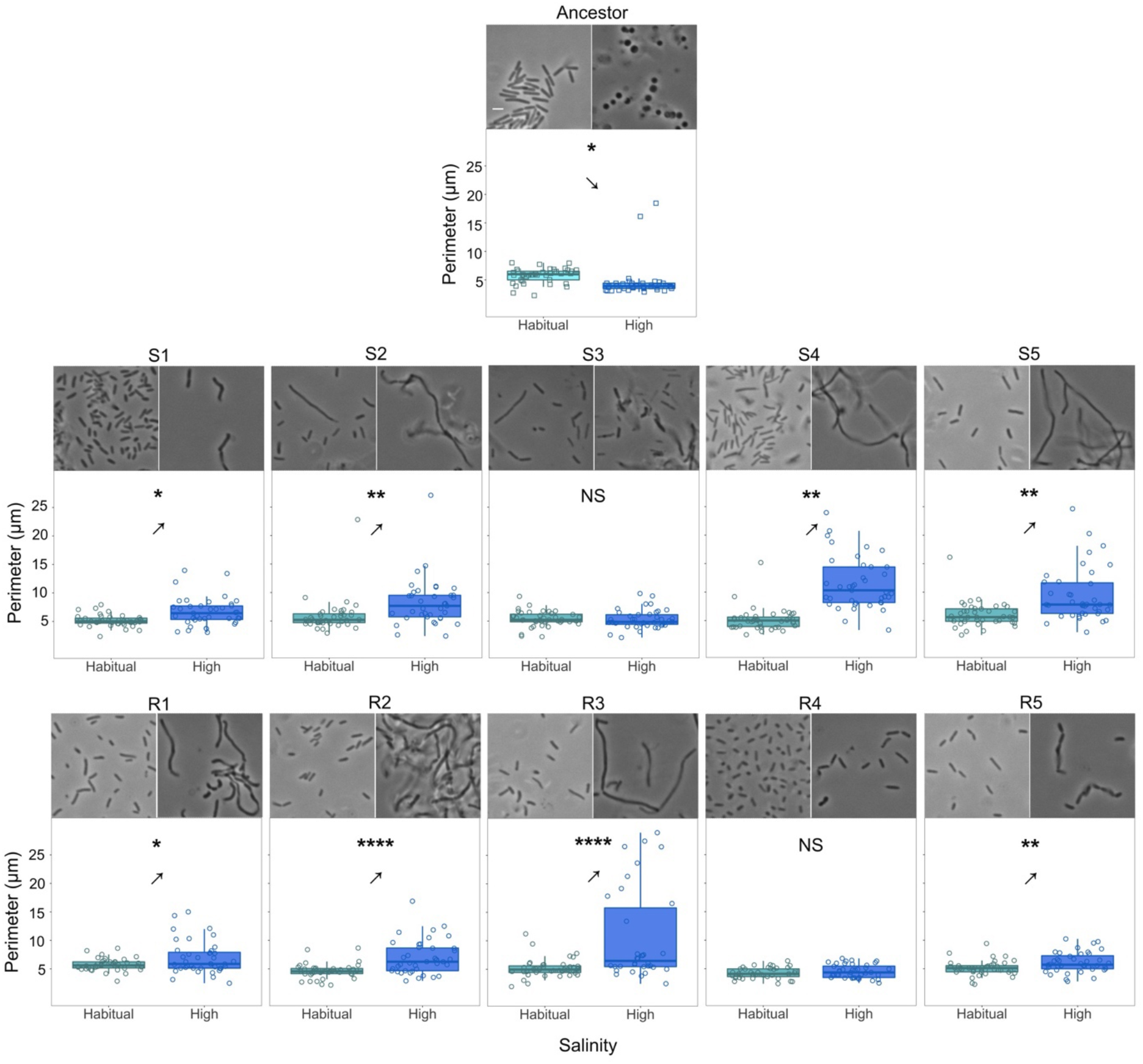
The evolution of phenotypic plasticity in cellular morphology. The arrows point towards the qualitative direction of phenotypic plasticity. *: *P* ≤ 0.05; **: *P* ≤ 0.01; ***: *P* ≤ 0.001; ****: *P* ≤ 0.0001. See Table S1 for statistical details. The reversal of the ancestral cell perimeter plasticity (observed in eight out of 10 evolved clones) corresponded to the canalization of phenotypically plastic cell clustering (compare with Fig. 2b). Also see Fig. S6 for cell shape plasticity quantified in terms of cellular circularity.

The ancestral genotype showed significant phenotypic plasticity in terms of the cellular perimeter observed in 2d images: specifically, the ancestor had significantly smaller cells under high salinity than under habitual salinity (Fig. 3; Table S1). In contrast, clones representing 4 out of five S lines (S1, S2, S4, and S5) and 4 out of five R lines (R1, R2, R3, and R5) showed a reversal of the ancestral phenotypic plasticity in terms of the cell perimeter (Fig. 3; Table S1). Specifically, S1, S2, S4, S5, R1, R2, R3, and R5 showed significantly larger cells under high salinity (Fig. 3; Table S1). There was a clear correspondence between reversal of the cell perimeter plasticity and successful canalization of cellular clustering: The eight lines that showed reversal in the ancestral cell perimeter plasticity were also the ones that successfully canalized the cellular clustering during experimental evolution (compare Figs. 2 and 3). On the other hand, the S3 and R4 clones showed no difference in cellular perimeters under habitual versus high salinity while also failing to successfully canalize cellular clustering (compare Figs. 2 and 3).

We also analyzed cellular morphology in terms of the circularity of individual cells (see *Methods*). The ancestor showed significant phenotypic plasticity in terms of cell circularity: (rod shaped cells under habitual salinity versus spherical cells under high salinity; Fig. S6; Table S2). In contrast, the cells belonging to S1, S3, R4, and R5 underwent moderate elongation that imparted the characteristic rod shape of *E. coli* under both habitual and high salinity (Fig. S6). Moreover, S1, S3, R1, R4, and R5 underwent canalisation in terms of their cellular circularity (*i.e.*, their cells exhibited similar circularity under both habitual and high salinity (Fig. S6; Table S2)). A relatively greater cellular elongation in S2, S4, S5, R2, and R3 under high salinity reversed their ancestral phenotypic plasticity in terms of cellular circularity and made them filamentous (Fig. 3; Table S1)).

Having investigated phenotypic plasticity and its canalization at two distinct levels of biological organization (selectable multicellular units vs. individual cells), we studied the genetic basis of the evolution of undifferentiated multicellularity observed in our experiments.

### The genetic basis of canalized multicellularity in *E. coli*

We sequenced whole genomes of all the S and R clones described in Figs. 2 and 3 and compared them to the ancestor to identify the mutations that resulted in the evolution of simple macroscopic multicellularity in our experiments (see *Methods*). We found that most mutations occurred within genes involved in the biosynthesis of peptidoglycan, which forms the bulk of eubacterial cell walls (Fig. 4a). Specifically, out of the 19 mutations putatively linked to changes in cell surface properties, 13 were found within genes directly involved in peptidoglycan biosynthesis (Table S3). We found that 7 out of the ten sequenced clones had a mutation in MraY, the enzyme that catalyzes the first membrane-bound step of peptidoglycan biosynthesis^42^. Despite such high degree of parallelism at the level of genes, we found several different mutations at widely distributed locations within the primary chain of MraY (Table S3). Interestingly, all these mutations were concentrated towards one side of the tertiary structure of MraY, facing the periplasmic zone of the transmembrane protein (Fig. 4b). These mutations, spaced apart from the cytoplasmic active site of MraY by the bacterial inner membrane, likely play a role in recruiting other peptidoglycan-related proteins in the periplasm. We found that all the seven clones with an MraY mutation successfully evolved simple macroscopic multicellularity by canalizing the ancestrally plastic cellular clustering (compare Figs. 2b and 4b). In addition to an MraY mutation, 6 of these seven clones also carried mutations in other genes linked to peptidoglycan synthesis, biofilm formation, or adaptation to nutrient media (Table S3). Moreover, we observed the strongest canalization of the clustering phenotype (with almost all bacterial growth within macroscopic clusters and the absence of detectable turbidity) in the S clones that carried at least one mutation in addition to an MraY mutation (S1, S2, and S5; see Fig. 2b and Table S2).

**Figure 4.**
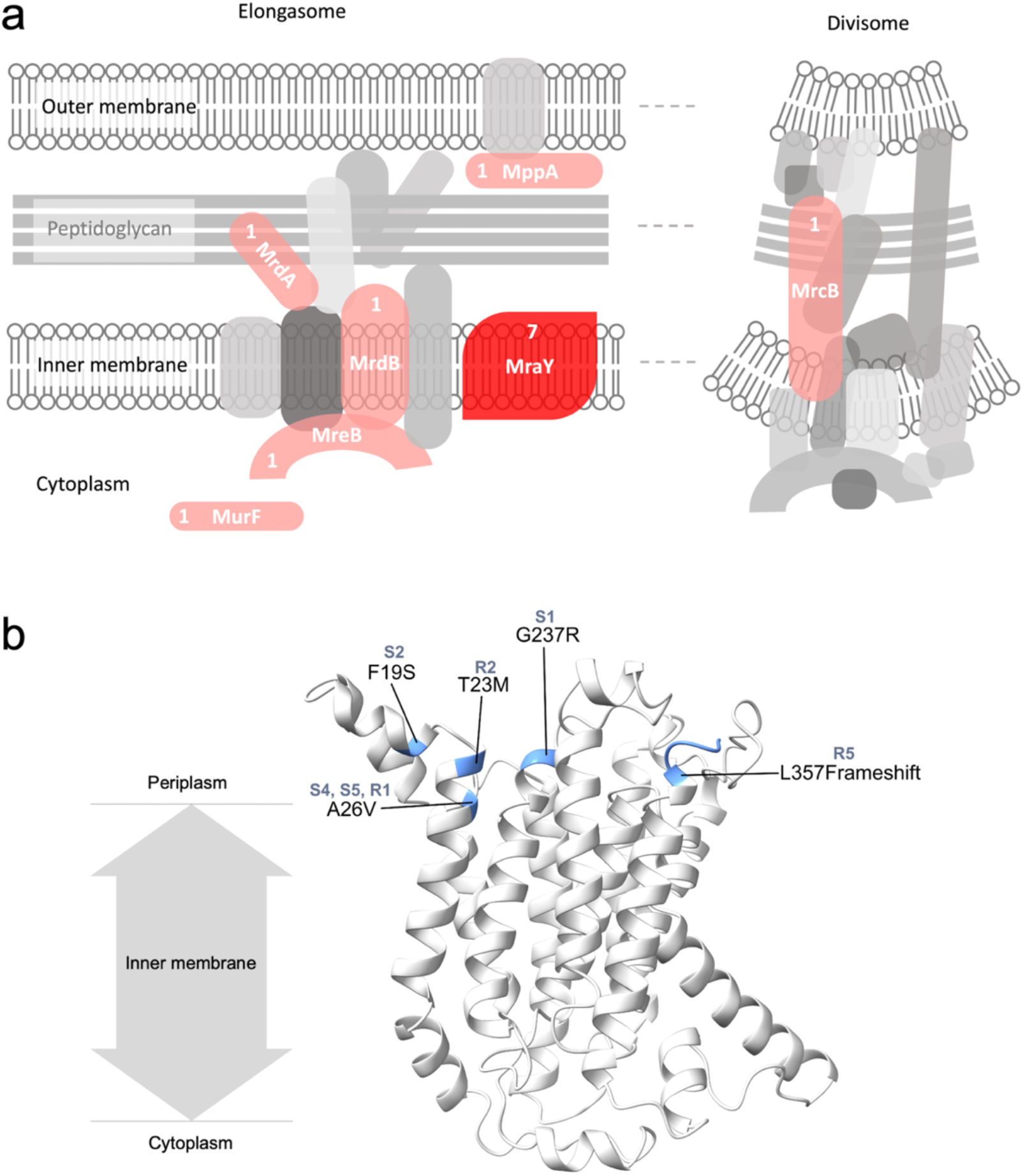
The genetic basis of canalized multicellularity. (a) Experimental evolution of canalized multicellularity primarily enriched mutations in genes involved in cell wall assembly. The schematic shows the proteins encoded by the mutated genes in red. The numbers accompanying the mutated proteins represent the number of clones that showed a mutation in a particular protein. Two genes (*murF* and *mppA*) showed synonymous mutations. (b) The location of mutations on the 3D structure of MraY, the protein that mutated in 70% of the sequenced clones. All the mutated regions are located near the periplasmic region of the transmembrane protein.

Curiously, S4 was only one MraY mutation away from the common ancestor (Table S3), which suggests that a single mutation can be sufficient for canalizing the ancestrally plastic cell clustering. It is worth noting that such canalization driven by a single mutation was relatively weak: the S4 clone showed a combination of macroscopic clusters and planktonic growth under habitual salinity (Fig. 2b). Furthermore, R3, the only clone that evolved a multicellular life history without enriching an MraY mutation, also displayed a weak canalization characterized by a combination of both clustering and turbid growth (Fig. 2b).

Unlike the S clones, the R clones preferentially colonized the air-liquid interface (Supplementary movies 5-8). We found that the mutations in the R clones could potentially explain this phenotypic difference. Specifically, R5 had two mutations in genes putatively linked to mat formation at the air-liquid interface through c-di-GMP signaling (*dgcQ* and *pdeA* (Table S3)). Moreover, R2, R3, and R4 had mutations within (or upstream to) genes with possible links to biofilm formation (*mprA* encoding a transcriptional repressor (R2, R3, and R4) and *bhsA* encoding an outer membrane protein (R4); Table S3). Furthermore, none of the S clones showed a mutation in any of these four genes linked to interface inhabiting mat formation (*dgcQ, pdeA, mprA, or bhsA*). Such mutational contrast could potentially explain the differences in the abilities of the S and R clones to inhabit the air-liquid interface.

Several other genes linked to peptidoglycan biosynthesis which mutated in our study (*mrcB, mrdA, mrdB, mreB, murF*; Fig. 4a) are linked to the maintenance of cell shape in *E. coli*^43–45^. Specifically, both MrdA and MrdB are known to play key roles in maintaining the characteristic rod shape of *E. coli*^43,44^. Moreover, MreB, which is the bacterial analogue of actin, is an essential protein that forms a scaffold which interacts with several other peptidoglycan biosynthesis proteins and plays key role in cellular elongation^42^. Finally, MraY has been shown to affect both the cell shape and adhesion in the multicellular cyanobacterium *Anabaena*^46^. This suggests that the mutations observed in the clones that successfully canalized multicellular clustering can also be linked to the evolutionary changes in cell shape plasticity (Fig. 3; Table S3).

## Discussion

Our study begins with the demonstration that bacteria show phenotypically plastic cell clustering that results in large macroscopic structures in high salinity environments. Since both Gram-negative and Gram-positive bacteria exhibit this phenomenon (Fig. 1), such plastic development of multicellular clusters appears to be a common (but not universal) bacterial capacity. Interestingly, the trigger for such phenotypically plastic cell clustering (high salinity) is frequently encountered by bacteria in diverse environments ranging from marine habitats to human skin. Therefore, such clustering is expected to have important ecological implications. We further showed that this plastic capacity to form multicellular clusters is evolvable and can be canalized rapidly to result in bacteria that obligately grow as multicellular clusters.

Our study is unique because it demonstrates not only that phenotypic plasticity can facilitate the evolution of macroscopic multicellularity, but also that it can do so in unicellular bacteria. Specifically, although previous studies have shown that the canalization of phenotypic plasticity can lead to multicellular development^11,31,47^, their multicellular structures contained < 200 cells and remained microscopic. Moreover, a recent important study has demonstrated the mutation-driven (*i.e.*, not plasticity-based) evolution of macroscopic multicellularity in a eukaryote (yeast), where the largest multicellular clusters comprised ~ 4.5 × 10^5^ cells^9^. Building on this fascinating finding, we show that phenotypic plasticity can enable unicellular *bacteria* to form macroscopic clusters comprising > 10^5^ CFUs under high salinity. Furthermore, we successfully canalized this plastic phenotype to form macroscopic clusters comprising > 10^4^ CFUs without any environmental induction. We also note that our CFU counts within clusters are likely underestimates (see *Methods*). Furthermore, since the evolved bacteria grow obligately as macroscopic multicellular clusters even in the absence environmental induction, we conclude that they have successfully evolved the first step towards the multicellularity that requires the obligate formation of undifferentiated clusters. Importantly, such obligately multicellular growth of our evolved bacteria is distinct from the facultative formation of largely planar biofilms (with limited vertical growth) upon attachment to substrate surfaces, as shown by diverse bacterial species^48^.

The evolution of multicellularity is considered to be one of the most frequent ‘major transitions’ because a large diversity of ecological conditions can make multicellularity selectively favorable^5,49^. Corroborating this notion, our results suggest that owing to phenotypically plasticity, the ability to evolve multicellularity should be widespread among bacteria, which comprise a rather large part of the tree of life. Crucially, plastic clustering enables bacteria to avoid waiting for the selection of specific *de novo* mutations that make cells stay together. Instead, environmental changes (*e.g.*, an increase in salinity) can rapidly lead to the development of multicellular phenotypes, which could then be subjected to selection. By demonstrating this ‘genes as followers’ mode of evolution^34^, our study also highlights the role of plasticity in a major evolutionary transition. Although most studies dealing with phenotypic plasticity tend to investigate one plastic trait^50^, some studies have led to powerful insights by simultaneously investigating plasticity in multiple traits, all of which belong to the same level of biological organization^51–53^. Our study makes a significant advance in this field by investigating phenotypic plasticity and its canalization at two different levels of biological organization (*collectives* of cells (Fig. 2b) and *individual* cells (Fig. 3)). An important aspect of our experiment is that it demonstrates the simultaneous evolution of plasticity in opposite directions at different levels of organization (compare Figs 2b and 3). Specifically, at the level of cell collectives, most of the evolved lines formed multicellular clusters under both habitual and high salinity; this phenotype was ancestrally expressed *in the presence of environmental induction* (Fig. 2b). In contrast, at the level of individual cells, the evolved lines showed non-spherical cell shapes with an average circularity of ≤ 0.667 under both low and high salinity; the ancestor expressed such cell circularity *in the absence of environmental induction* (Fig. 3). Taken together, these observations caution against forecasting an evolutionary change in phenotypes by extrapolating from the phenotypic plasticity shown by the ancestor.

Although both spherical and rod-shaped cells can form multicellular clusters under high salinity, our selection for greater clustering under progressively reducing environmental induction ended up selecting for elongated bacterial cells (Fig. 3). Moreover, we found that all the six clones that showed highly elongated (filamentous) cells under high salinity (S2, S4, S5, R1, R2, and R3) also exhibited efficient canalization of the multicellular clustering (Fig. 2b). On the other hand, the two clones which could not canalize multicellular clustering successfully (S3 and R4) also lacked highly elongated cells under high salinity (Fig. 3). Thus, cellular elongation under high salinity closely corresponded with the canalization of the ancestrally plastic cell clustering. This notion aligns with two recent eukaryotic studies which argue that greater cell elongation leads to more efficient packing within clusters^9,54^. It may also explain why the canalization of multicellularity was based on mutations predominantly in cell shape modulating peptidoglycan biosynthesis loci (Fig. 4). The highly parallel molecular evolution we observed at the level of loci points towards a putative pleiotropy between cell shape and clustering. This notion is strengthened by our observation that in clone S4, a single MraY mutation could not only canalize the ancestrally plastic cell clustering but also give rise to highly elongated cells (Figs. 2b and 3). Moreover, despite superficially resembling *Pseudomonas fluorescens* mats formed under static conditions, the mats formed by R1, R2, R3, and R5 under optimal salinity were genotypically different: Unlike *P. fluorescens* mats that are predominantly formed by mutants overproducing cyclic-di-GMP^55^, all our *E. coli* clusters were primarily caused by mutations in peptidoglycan biosynthesis genes (Figs. 2 and 4). Interestingly, in addition to an MraY mutation (linked to peptidoglycan biosynthesis), R5 also contained two mutations linked to cyclic-di-GMP expression (Table S3).

Apart from adding multiple key insights to the current understanding of how multicellularity evolves, our results should also act as stepping-stones for new theoretical and empirical studies in several diverse fields of inquiry (Fig. S8). For example, why bacteria tend to form a single columnar cluster under high salinity instead of multiple globular clusters is a fascinating biophysical puzzle. Moreover, a generic tendency to form environmentally induced clusters could significantly impact the ecological interactions between multiple different bacterial species, potentially facilitating long-term co-existence by providing spatially segregated growth. Furthermore, the cells at the cluster’s periphery inevitably face a different environment as compared to those at the core. Hence, an exciting new line of work would be to test if such ecological differences can drive the evolution of cellular differentiation. Finally, by demonstrating that bacteria can rapidly evolve macroscopic multicellularity, our results call for a reconsideration of why multicellular organisms are predominantly eukaryotic.

## Methods

### Bacterial strains and nutrient media

We used the following bacteria for studying the phenotypic plasticity of cell clustering: *Escherichia coli* K12 substr. MG1655 (Eco galK::cat-J23101-dTomato); *Staphylococcus aureus* JE2; *Pseudomonas aeruginosa* PAO1; *Citrobacter freundii* ATCC 8090; *Serratia marcescens* BS 303. The bacteria were cultured in liquid environments containing Luria Bertani broth (10 g/L tryptone, 5 g/L yeast extract) with 5 g/L NaCl (habitual salinity) or 60 g/L NaCl (high salinity).

### Timelapse movies

We used Canon Rebel T3i (Canon Inc. (Ōta, Tokyo, Japan)) to capture macroscopic images and then stitched them into timelapse movies using Persecond for Mac version 1.5 (Flixel Inc. (Toronto, Canada)). For all the timelapses reported in our study, we used a remote control to automatically capture an image every 4 minutes and published the movie files at 16 fps.

### Experimental evolution

We conducted experimental evolution with bacterial populations derived clonally from a single *E. coli* MG1655. We propagated five independent replicate populations each belonging to two distinct selection lines (S (for Shaken (at ~180 rpm)) and R (for Resting)) by culturing bacteria in glass tubes containing 5 ml LB (Fig. 2a). In the beginning of the evolution experiment, the bacteria were cultured in Luria Bertani broth supplemented with 6% NaCl (w/vol). The NaCl concentration in the nutrient medium was progressively reduced over 50 days during the experiment (6% w/vol (days 1-11), 5% w/vol (days 12-15), 4% w/vol (days 16-19); and 3% w/vol (days 20-50)). We subcultured bacteria into fresh nutrient medium every 24 h using a selection protocol designed to enrich cell clustering phenotypes in the face of progressively reducing environmental induction. For each subculture, we picked a small piece of the previous day’s bacterial cluster fitting within 20 μl and washed it serially in 2 ml fresh media in four distinct wells. This diluted the planktonically growing bacteria by 10^-8^-fold while keeping the clustered bacteria undiluted. We stored periodic cryo-stocks for all the 10 independently evolving populations. We streaked the endpoint cryo-stocks on Luria agar without any externally supplemented NaCl and isolated a colony from each population after 18 hours. We used these colonies (clones S1, S2, S3, S4, S5, R1, R2, R3, R4, and R5) to conduct growth assays and genomic sequencing.

### Microscopy and cell shape analysis

We performed both brightfield and fluorescence microscopy with clonal ancestral and evolved samples at 100x magnification (oil immersion) using Nikon Eclipse 90i (Nikon Inc. (Amstelveen, NL)). All the samples subjected to microscopy were streaked on fresh Luria agar from their respective cryo-stocks. A single colony was then used to inoculate the liquid media in question (high versus habitual salinity) to obtain the phenotype at the level of cell collectives. 5 μl samples from fully grown liquid cultures (containing planktonic cells and/or macroscopic clusters) were spotted on a glass slide and protected with a glass coverslip, which resulted in the flattening and disintegration of the clusters. We used the Texas Red optical filter (excitation: 562/40 nm; emission: 624/40 nm) to observe cells expressing dTomato. Overlays between brightfield and fluorescent images were used to identify cell shapes and boundaries. We used the open-source software FIJI (ImageJ 1.53) for Mac to analyze cell shapes by manually tracing the cellular boundaries. We computed cellular perimeter and circularity 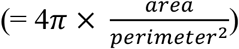 using built-in functions in FIJI.

### Statistics

#### Cell shape plasticity

We used two-tailed *t*-tests (unequal variance across types) to analyze the difference between cell shape parameters for a given genotype across habitual versus high salinity (N = 40 cells). The two cell shape parameters (perimeter and circularity) were analyzed separately.

#### CFU counts

We used single simple *t*-tests to compare the CFU counts of the evolved clones against the ancestral level, both under high and habitual salinity (N = 5 distinct clones each for S (S1, S2, S3, S4, and S5) and R (R1, R2, R3, R4, and R5)).

### Whole genome sequencing

Genomic DNA from single colonies from each population was isolated using GeneJet Genomic DNA Purification kit (Thermo Scientific™) for whole genome sequencing on the evolved clones and the ancestor. We used a standard miniaturized protocol to prepare DNA libraries using the NEBNext Ultra II FS DNA Library Prep Kit for Illumina (New England BioLabs Inc. (Ipswich, MA, USA))^56^. The quantity of the prepared DNA libraries was validated with a Qubit© 2.0 Fluorometer (Thermo Fisher Scientific Inc. (Waltham, MA, USA)). We used the MiSeq system (Illumina Inc. San Diego, CA, USA) to perform 250-bp paired end next generation sequencing on the prepared libraries at a minimum coverage of 10x (the average coverage of the detected mutations was 43.80x). We analyzed the sequencing output using the Geneious Prime software for Mac (v2022.0.2) and trimmed the sequencing output data using BBDuk to remove reads < 20 bp or with a quality score < 20. Since we conducted sequencing on clones, to avoid interpreting sequencing errors as mutations, we restricted our analysis to variants with frequencies ≥ 70%.

### Locating mutations on 3D protein structures

Since the crystal structure of MraY is not yet known for *E. coli*, a publicly available homology model made by Alphafold v2.0^57^ was used (accession P0A6W3). We used the UCSF ChimeraX software^58^ for Mac (https://www.rbvi.ucsf.edu/chimerax/) to identify and highlight the locations of the sites mutated in MraY in our experiment. The highlighted output was used to make Fig. 4b.

## Author contributions

Conceived the original idea and designed the project: Y.C.

Supervised the project: P.A.L.

Conducted the experiments and data analysis: Y.C.

Wrote the manuscript: Y.C. and P.A.L.

Acquired funding: Y.C. and P.A.L.

Refined the idea and provided key critiques: S.D.

## Acknowledgements

We thank Eric Libby, Jennifer Pentz, Anthony Sun, and Shraddha Karve for valuable discussions and constructive critiques. Y.C. was supported by a postdoctoral fellowship awarded by the Wenner-Gren Foundations (Sweden): Grants UPD2020-0113, UPD2021-0182. The funders had no role in study design, data collection and analysis, decision to publish, or preparation of the manuscript.

## Data availability

All relevant data are within the manuscript and its Supplementary Information files. The whole genome sequences reported in this study are available from the NCBI database (accession number: PRJNA880543; https://www.ncbi.nlm.nih.gov/sra/PRJNA880543).

