## Supplementary Information for "Bacteria evolve macroscopic multicellularity via the canalization of phenotypically plastic cell clustering"

*for*

Affiliations
[1]Department of Molecular Biology, Umeå University, Umeå, Sweden
[2]Indian Institute of Science Education and Research (IISER) Pune, Pune, India

### A note on the organization of *Supplementary files*

There are ten *Supplementary files* linked to the main manuscript (1 *.docx* file (the current document) and 9 *.mp4 Supplementary Movie* files). The *Supplementary Movie* files are mentioned and described here in the order in which they are referred to in the main manuscript. The *Supplementary Figures* are embedded here directly and are not presented as separate files.

**Supplementary movie 1**

File: Supplementary_movie_1.mp4

Description: A timelapse of the growth of the *E. coli* ancestral (wildtype) genotype under habitual salinity

Screenshot:

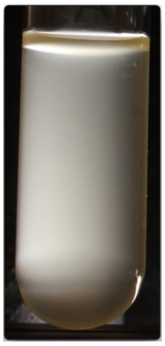

Frame rate: 16 fps
Duration: 20 s

**Supplementary movie 2**

File: Supplementary_movie_2.mp4

Description: A timelapse of the growth of *S. aureus* under habitual salinity

Screenshot:

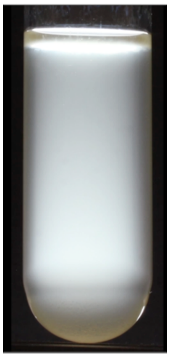

Frame rate: 16 fps
Duration: 16 s

**Supplementary movie 3**

File: Supplementary_movie_3.mp4

Description: A timelapse video of the growth of the *E. coli* ancestral (wildtype) genotype under high salinity

Screenshot:

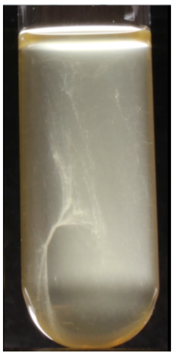

Frame rate: 16 fps
Duration: 20 s

**Supplementary movie 4**

File: Supplementary_movie_4.mp4

Description: A timelapse video of the growth of *S. aureus* under high salinity

Screenshot:

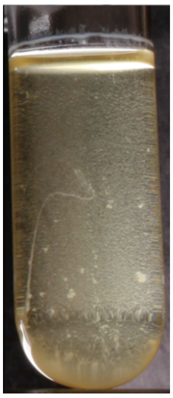

Frame rate: 16 fps
Duration: 16 s

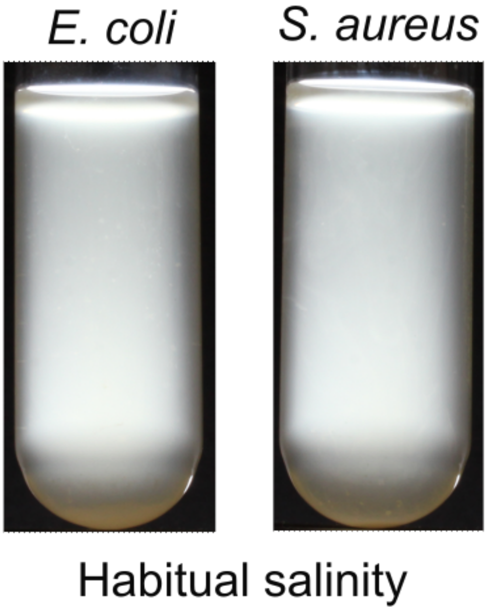


**Fig. S1. Both *E. coli* and *S. aureus* clusters disintegrate and grow planktonically upon being transferred to the habitual salinity environment (incubation period: 24 h).**

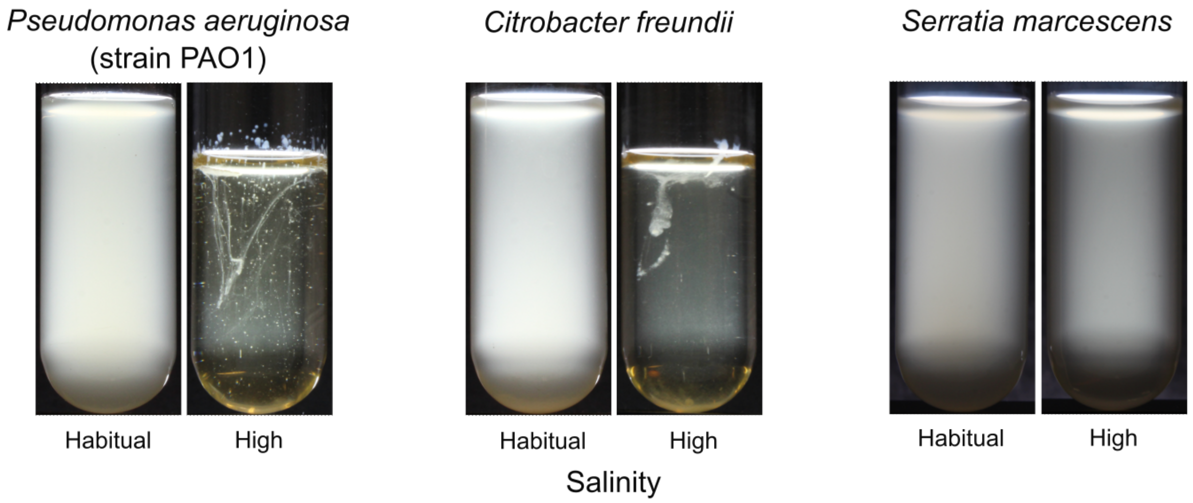


**Fig. S2. Phenotypic plasticity of bacterial growth under habitual versus high salinity.** Incubation times: *Pseudomonas aeruginosa* (strain PAO1): 24 h under habitual salinity, 72 h under high salinity (reduction in volume due to evaporation); *Citrobacter freundii*: 24 h under habitual salinity, 72 h under high salinity (reduction in volume due to evaporation); *Serratia marcescens*: 24 h under both habitual and high salinity.

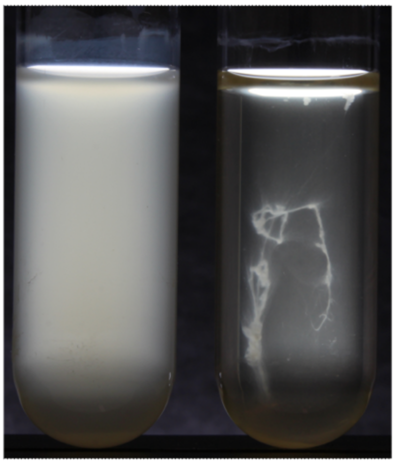


**Fig. S3 Phenotypically plastic macroscopic cell clustering exhibited by *E. coli* MG1655 under well mixed conditions in tubes shaken at ~180 rpm (incubation period: 24 h).**

**Supplementary movie 5**

File: Supplementary_movie_5.mp4

Description: A timelapse video of the growth of the S clones under habitual salinity. Order from left to right: S1, S2, S3, S4, and S5

Screenshot:

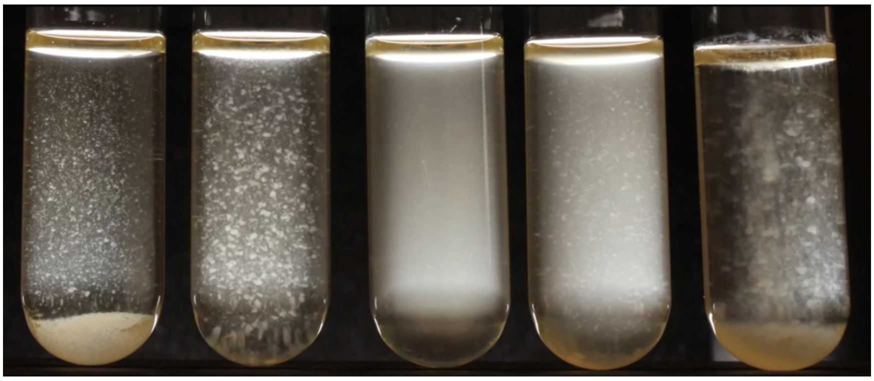

Frame rate: 16 fps
Duration: 20 s

**Supplementary movie 6**

File: Supplementary_movie_6.mp4

Description: A timelapse video of the growth of the S clones under habitual salinity. Order from left to right: R1, R2, R3, R4, and R5

Screenshot:

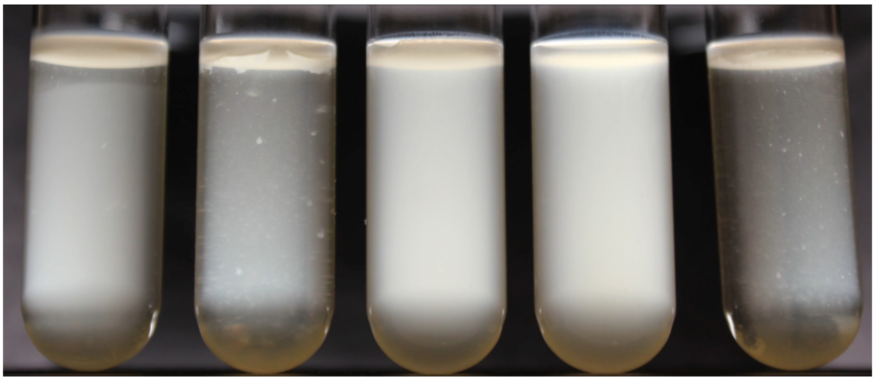

Frame rate: 16 fps
Duration: 19 s

**Supplementary movie 7**

File: Supplementary_movie_7.mp4

Description: A timelapse video of the growth of the S clones under high salinity. Order from left to right: S1, S2, S3, S4, and S5

Screenshot:

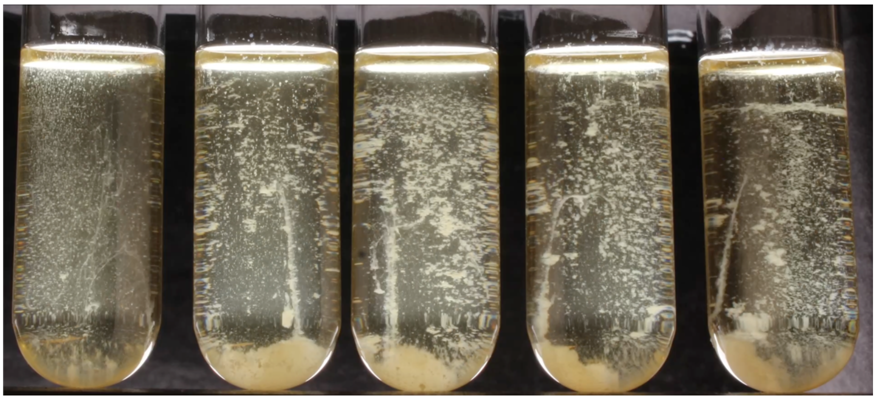

Frame rate: 16 fps
Duration: 34 s

**Supplementary movie 8**

File: Supplementary_movie_8.mp4

Description: A timelapse video of the growth of the R clones under high salinity. Order from left to right: R1, R2, R3, R4, and R5

Screenshot:

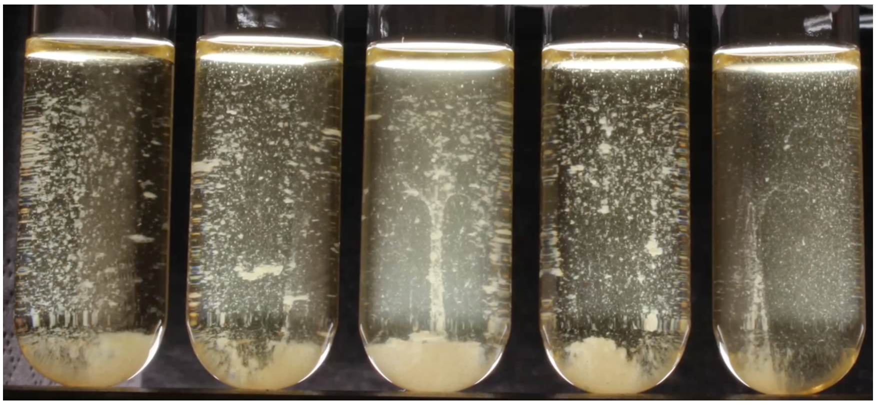

Frame rate: 16 fps
Duration: 34 s

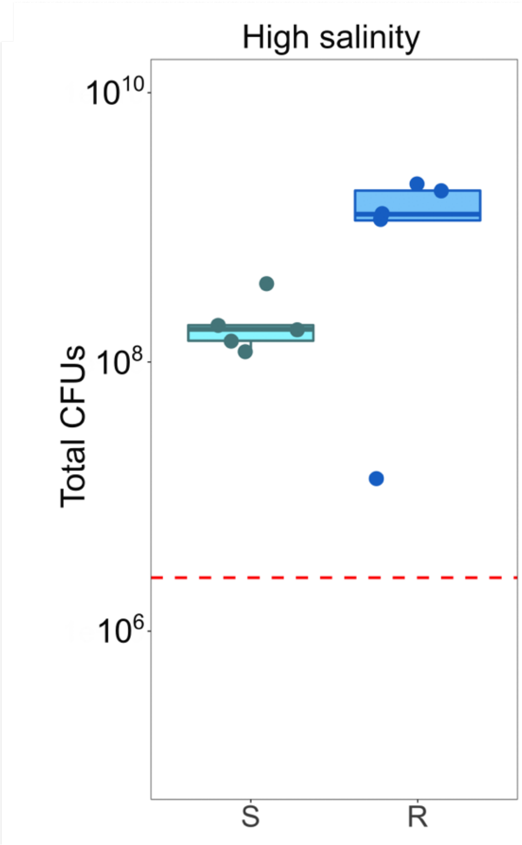


**Fig. S4. The total number of colony forming units (CFUs) detected after 24 hours of growth under high salinity in 5 ml medium.** The red dashed line represents the ancestral level. The CFU counts are likely underestimates due to the possibility of incomplete cluster breakage upon vortexing prior to CFU determination. See *Methods* for details.

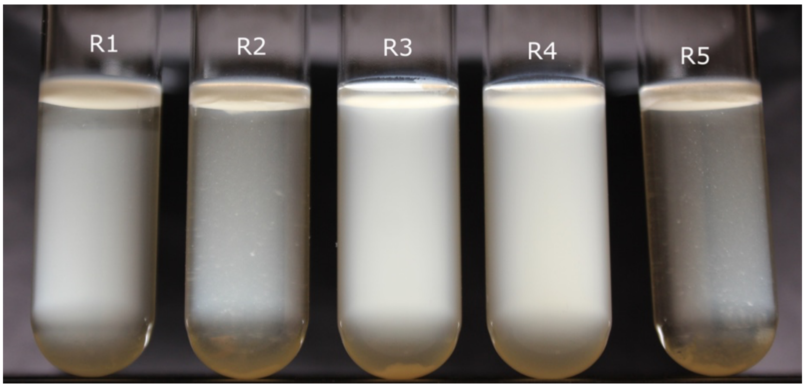


**Fig. S5. The R1-R5 clones grown under habitual salinity pictured just prior to external perturbation. R1, R2, R3, and R5 successfully canalized macroscopic multicellularity by inherently growing as interface dwelling mats even without environmental induction. Also see Fig. 2b.**

**Supplementary movie 9**

File: Supplementary_movie_9.mp4

Description: A timelapse video depicting that old clusters can spontaneously give rise to new clusters upon accessing fresh medium. Left: S5 under habitual salinity; right: S5 under high salinity.

Screenshot:

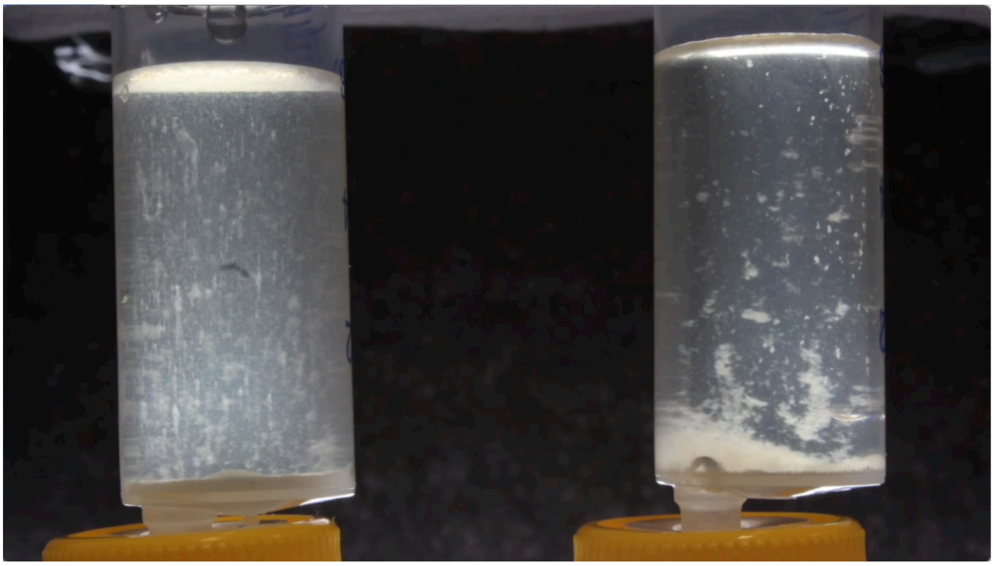

Frame rate: 16 fps

Duration: 25 s

**Table S1. Summary of two-tailed *t*-tests (unequal variance across types) on cellular perimeter across habitual and high salinity (N = 40).**

| Clone | Mean perimeter under habitual salinity (μm) | Mean perimeter under high salinity (μm) | *P* value |
|---|---|---|---|
| Anc | 5.78 | 4.529 | 0.02 |
| S1 | 5.132 | 7.508 | 0.011 |
| S2 | 5.887 | 10.114 | 0.002 |
| S3 | 5.385 | 5.323 | 0.854 |
| S4 | 5.209 | 14.711 | 0.007 |
| S5 | 6.048 | 10.426 | 0.002 |
| R1 | 5.747 | 6.949 | 0.017 |
| R2 | 4.618 | 6.964 | $< 10^{-5}$ |
| R3 | 5.174 | 15.078 | $< 10^{-5}$ |
| R4 | 4.236 | 4.522 | 0.226 |
| R5 | 5.134 | 6.094 | 0.008 |

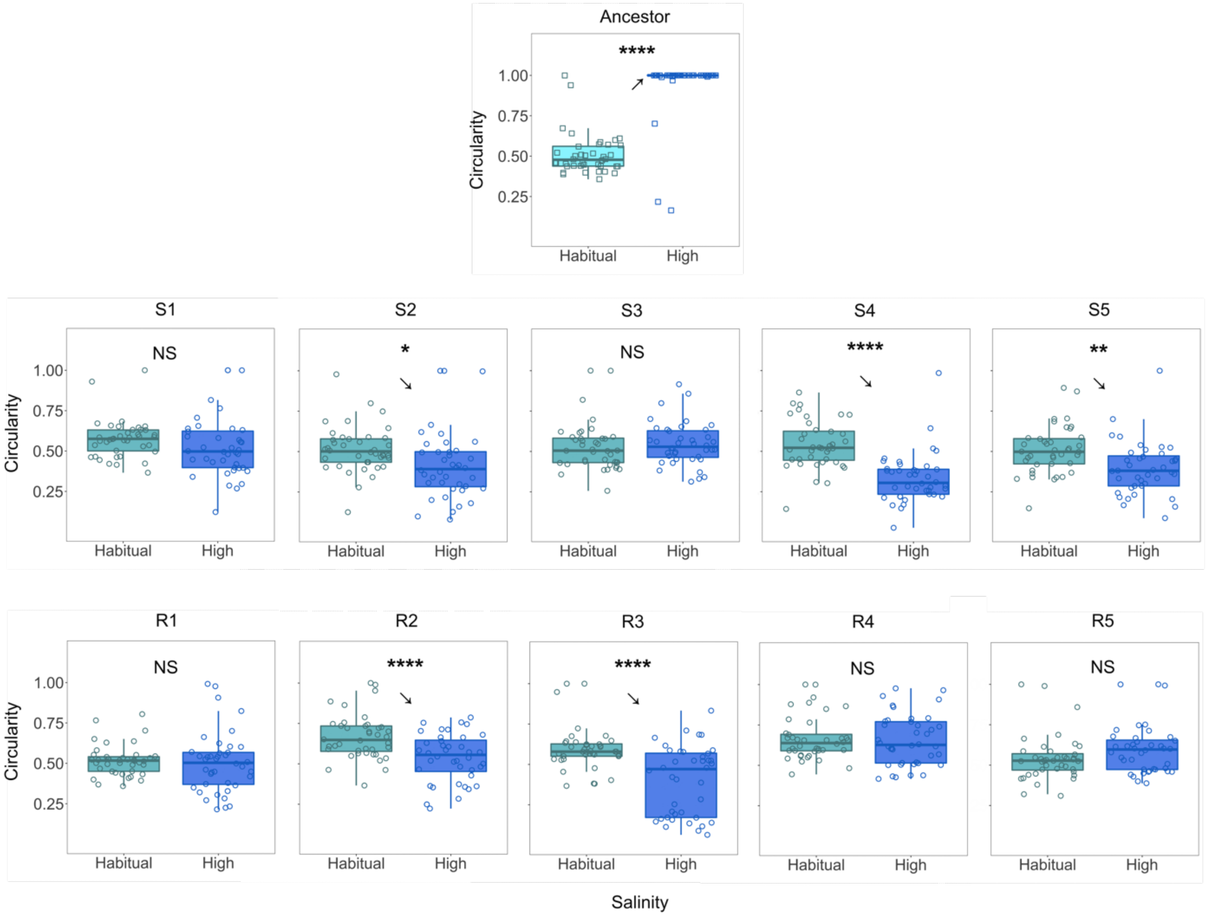


**Figure S6. The evolution of phenotypic plasticity in cellular circularity.** The arrows point towards the qualitative direction of phenotypic plasticity. *: $P \leq 0.05$; **: $P \leq 0.01$; ***: $P \leq 0.001$; ****: $P \leq 0.0001$. See Table S1 for statistical details.

**Table S2. Summary of two-tailed *t*-tests (unequal variance across types) on cellular circularity across habitual and high salinity (N = 40).**

| Clone | Mean circularity under habitual salinity | Mean circularity under high salinity | *P* value |
|---|---|---|---|
| Anc | 0.51 | 0.951 | $< 10^{-19}$ |
| S1 | 0.576 | 0.528 | 0.19 |
| S2 | 0.514 | 0.419 | 0.026 |
| S3 | 0.53 | 0.545 | 0.629 |
| S4 | 0.54 | 0.334 | $< 10^{-8}$ |
| S5 | 0.508 | 0.393 | 0.002 |
| R1 | 0.518 | 0.504 | 0.673 |
| R2 | 0.667 | 0.538 | $< 10^{-4}$ |
| R3 | 0.598 | 0.4 | $< 10^{-6}$ |
| R4 | 0.666 | 0.655 | 0.731 |
| R5 | 0.543 | 0.597 | 0.11 |

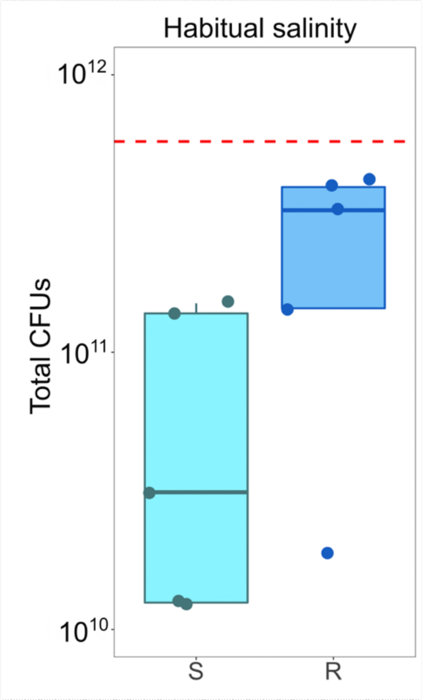


**Fig. S7. The total number of colony forming units (CFUs) detected after 24 hours of growth under habitual salinity in 5 ml medium.** The red dashed line represents the ancestral level. Both S and R CFU counts (but not the ancestral level) are likely underestimates due to the possibility of incomplete cluster breakage upon vortexing prior to CFU determination. See *Methods* for details. Single sample *t*-tests against the ancestral value (N = 5): $P < 10^{-4}$ (S clones); $P = 0.0151$ (R clones).

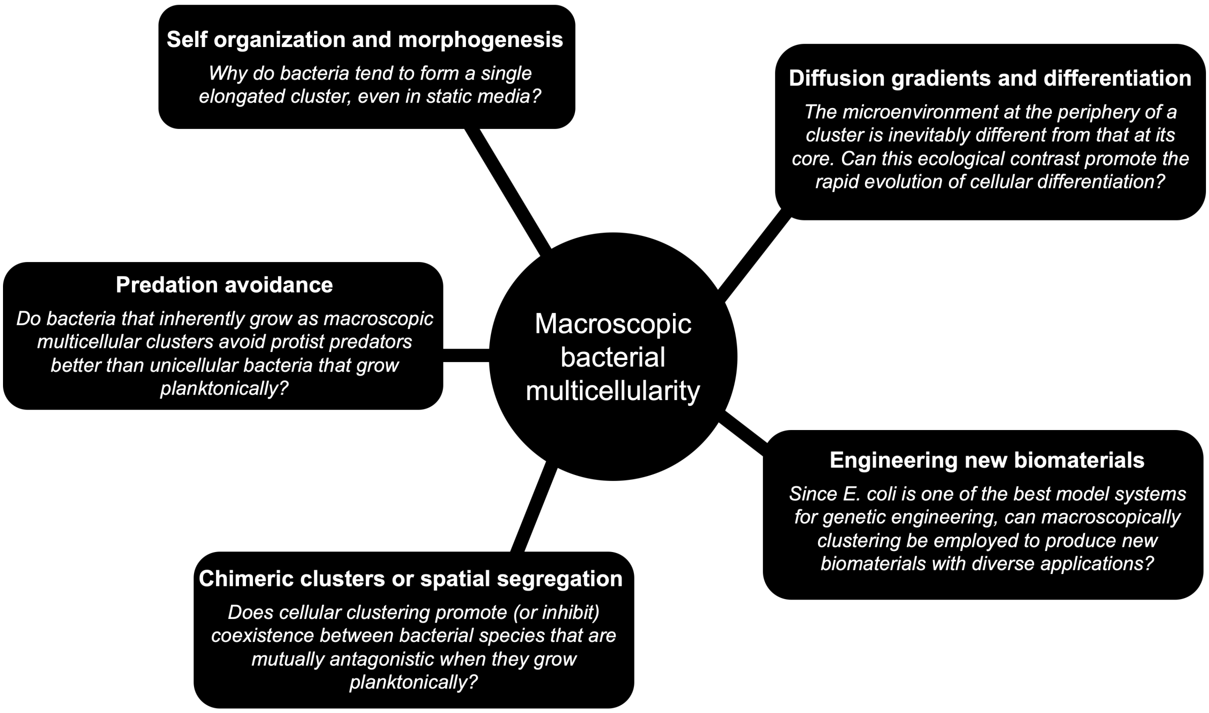


**Fig. S8. Some important future research questions and directions likely to spring forth from our results.** Taken together, our results should be of interest to a wide variety of scientific fields.
